# Unsupervised outlier detection applied to SARS-CoV-2 nucleotide sequences can identify sequences of common variants and other variants of interest

**DOI:** 10.1101/2022.05.16.492178

**Authors:** Georg Hahn, Sanghun Lee, Dmitry Prokopenko, Jonathan Abraham, Tanya Novak, Julian Hecker, Michael Cho, Surender Khurana, Lindsey R. Baden, Adrienne G. Randolph, Scott T. Weiss, Christoph Lange

## Abstract

As of June 2022, the GISAID database contains more than one million SARS-CoV-2 genomes, including several thousand nucleotide sequences for the most common variants such as delta or omicron. These SARS-CoV-2 strains have been collected from patients around the world since the beginning of the pandemic. We start by assessing the similarity of all pairs of nucleotide sequences using the Jaccard index and principal component analysis. As shown previously in the literature, an unsupervised cluster analysis applied to the SARS-CoV-2 genomes results in clusters of sequences according to certain characteristics such as their strain or their clade. Importantly, we observe that nucleotide sequences of common variants are often outliers in clusters of sequences stemming from variants identified earlier on during the pandemic. Motivated by this finding, we are interested in applying outlier detection to nucleotide sequences. We demonstrate that nucleotide sequences of common variants (such as alpha, delta, or omicron) can be identified solely based on a statistical outlier criterion. We argue that outlier detection might be a useful surveillance tool to identify emerging variants in real time as the pandemic progresses.

## 1. Introduction

More than one million nucleotide sequences of the SARS-CoV-2 virus have been collected from patients around the world since the beginning of the pandemic and made available in the GISAID database (Elbe and Buckland-Merrett, 2017; Shu and McCauley, 2017). Among them are more thousands of nucleotide sequences of the most common variants, precisely for the alpha (B.1.1.7), beta (B.1.351), delta (B.1.617.2), gamma (P.1), GH (B.1.640), lambda (C.37), mu (B.1.621), and omicron (B.1.1.529) variants (UCSC Genome Browser, 2022).

The emergence of new variants of the SARS-CoV-2 virus poses a threat to the progress made by ongoing vaccination campaigns against COVID-19 worldwide. Therefore, the detection and possible identification of newly emerging variants of the SARS-CoV-2 virus in (close to) real time is therefore of great interest.

Currently, a tool called “genomic surveillance” is used by the Centers for Disease Control (CDC) to detect new variants (CDC 2022a). This is done both through the National SARS-CoV-2 Strain Surveillance (NS3) program, as well as through commercial and academic laboratories contracted by the CDC, where genetic information of SARS-CoV-2 specimen are analyzed and classified into variants. By definition, a variant is characterized by having one or more mutations which differentiate it from other variants of the SARS-CoV-2 virus (CDC 2022b). A group of variants with similar genetic changes (a lineage) can be classified as a variant of concern (VOC) or a variant of interest (VOI) if they share characteristics that potentially necessitate public health action. For example, the U.S. government SARS-CoV-2 Interagency Group (SIG) classified omicron as a Variant of Concern (VOC) on 30 November 2021 due to the fact that omicron emerged in multiple countries without apparent travel history, the replacement of certain delta variants as predominant variants in South Africa by omicron, and its number of mutations in the spike protein which indicated a reduced susceptibility to sera from vaccinated individuals and certain monoclonal antibody treatments. The purpose of this article is to explore the ability of new unsupervised learning methodology that can to detect new variants of interest.

As shown previously in the literature (Hahn et al., 2020a,b), an unsupervised cluster analysis in which the similarity of all pairs of nucleotide sequences is assessed using the Jaccard index, and subsequent application of principal component analysis to the Jaccard similarity matrix, results in clusters of sequences according to certain characteristics such as their strain or their clade. Importantly, Hahn et al. (2020f) notice that nucleotide sequences the omicron variant cluster among sequences stemming from variants identified earlier on during the pandemic.

This finding immediately prompts the question whether the nucleotide sequences belonging to common variants can be identified by unsupervised outlier detection. In this article, we investigate this question by applying outlier detection to nucleotide sequences, both before the emergence of a variant and after a variant has emerged. We demonstrate that indeed, the number of detected outliers often increases shortly after the emergence of a new variant, and that nucleotide sequences of common variants can be identified solely based on a statistical outlier criterion.

Our findings could have important implications for the automated unsupervised identifications of SARS-CoV-2 strains. We argue that outlier detection might be a useful surveillance tool to identify emerging variants of interest in real time as the pandemic progresses.

The article is structured as follows. Section 2 introduces the methodology we use for this article, starting with data acquisition and cleaning, and how the similarity of sequences is assessed. We then describe the outlier detection method we use. Section 3 presents our findings on the clustering and outlier detection of SARS-CoV-2 nucleotide sequences. The article concludes with a discussion in Section 4.

## 2. Methods

In this section, we highlight methodological features of the analysis. In particular, we describe data acquisition and cleaning (Section 2.1), the assessment of the similarity of nucleotide sequences (Section 2.2), the methods used for outlier detection among sequences (Section 2.3), and the calibration of the outlier detection (Section 2.4).

### 2.1 Data acquisition and cleaning

All findings reported in this article are based on an image of all available SARS-CoV-2 nucleotide sequences on the GISAID database (Elbe and Buckland-Merrett, 2017; Shu and McCauley, 2017) from 28 March 2022, consisting of 211,167 sequences having accession numbers in the range of EPI_ISL_403962 to EPI_ISL_11498019. Sequences are only included in the analysis if they satisfy the four data quality attributes on GISAID. To be precise, all nucleotide sequences have to satisfy the criterion of being *complete* (defined as sequences having length at least 29,000bp), *high coverage* (defined as sequences with less than 1% N-bases), *with patient status* (defined as submissions with meta information consisting of age, sex, and patient status), and *collection data complete* (defined as submissions with a complete year-month-day collection date).

We aim to investigate if it is possible to detect sequences of a new variant among the other sequences in circulation upon emergence of that new variant. We consider eight common SARS-CoV-2 variants available on GISAID. Those are alpha (B.1.1.7), beta (B.1.351), delta (B.1.617.2), gamma (P.1), GH (B.1.640), lambda (C.37), mu (B.1.621), and omicron (B.1.1.529) variants.

To detect a new variant, we generate two reference datasets for each variant. For the first dataset, we determine the timepoint T1 at which the first sequences of each variant under consideration emerge on GISAID. We then generate the first reference dataset using only sequences from GISAID with a timestamp before T1. The second dataset emulates the emergence of a new variant. For this we determine the timepoint T2 at which 10% of all the sequences of a variant under consideration are available on GISAID (the threshold of 10% is arbitrary). We then generate the second reference dataset using only sequences from GISAID with a timestamp up to T2. The details of the reference datasets (the total number of sequences, their accession numbers on GISAID, as well as the time period they cover), are given in Tables 2 and 3.

**Table 1.** Local outlier detection approach. Number of detected outliers in Figures 5–12 before and after the emergence of each of the eight variants. True positives among the detected outliers, and number of sequences included for each variant.

| Variant | Before emergence of variant |  |  | After emergence of variant |  |  |
| --- | --- | --- | --- | --- | --- | --- |
|  | No. outliers | true positives | No. seq. | No. outliers | true positives | No. seq. |
| alpha | 1314 | 0 | 0 | 1070 | 329 | 788 |
| beta | 78 | 0 | 0 | 1902 | 88 | 99 |
| delta | 0 | 0 | 128 | 212 | 175 | 1085 |
| gamma | 1589 | 0 | 0 | 97 | 3 | 140 |
| GH | 137 | 0 | 0 | 179 | 0 | 4 |
| lambda | 2067 | 0 | 0 | 2066 | 4 | 4 |
| mu | 0 | 0 | 0 | 0 | 0 | 16 |
| omicron | 191 | 0 | 0 | 276 | 19 | 25 |

**Table 2.**
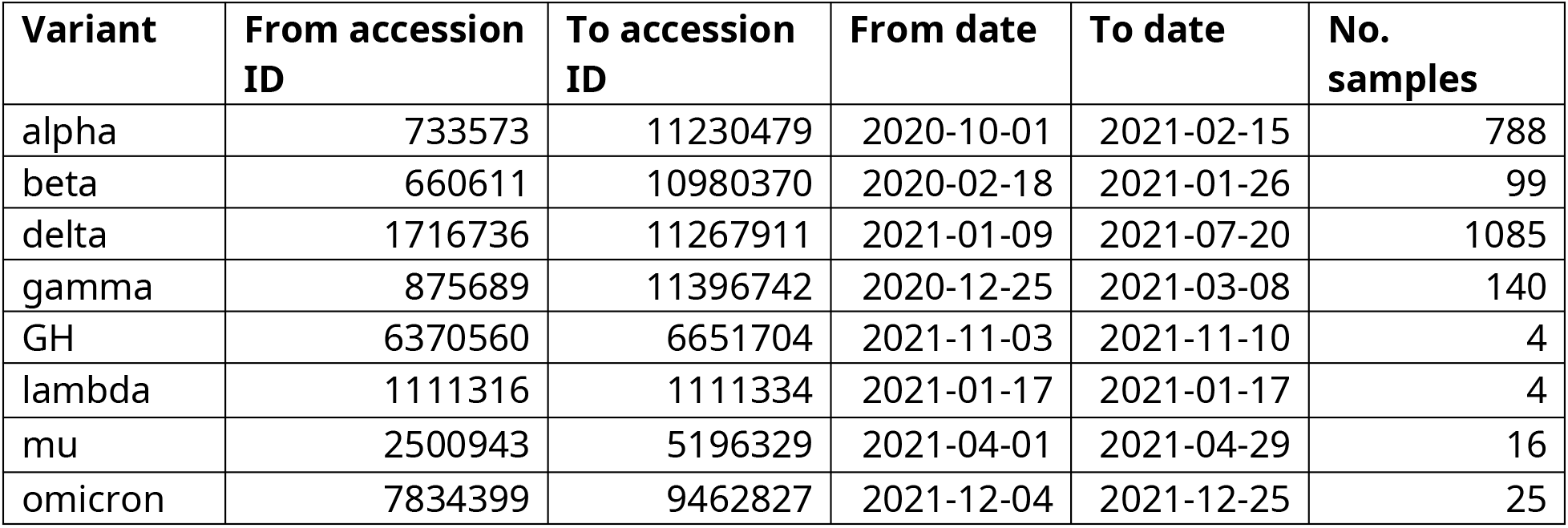
Composition of the dataset we aim to detect by variant. Range of accession numbers extracted from the GISAID database, their time stamps, and the total number of sequences included.

**Table 3.** Composition of the reference dataset we use to detect the emergence of a new variant. Range of accession numbers extracted from the GISAID database, their time stamps, and the total number of sequences included.

| **Variant** | **From accession ID** | **To accession ID** | **From date** | **To date** | **No. samples** |
| --- | --- | --- | --- | --- | --- |
| alpha | 406592 | 11403614 | 2020-01-08 | 2021-02-15 | 10000 |
| beta | 403963 | 11229964 | 2020-01-10 | 2021-01-26 | 9999 |
| delta | 404227 | 11403612 | 2020-01-16 | 2021-07-20 | 9999 |
| gamma | 407079 | 11448683 | 2020-01-10 | 2021-03-08 | 10000 |
| GH | 404227 | 11468151 | 2020-01-10 | 2021-11-22 | 10000 |
| lambda | 406593 | 11330894 | 2020-01-10 | 2021-01-17 | 9999 |
| mu | 408489 | 11468147 | 2020-01-10 | 2021-04-29 | 10000 |
| omicron | 410301 | 11448664 | 2020-01-13 | 2021-12-25 | 10000 |

Our planned subsequent computations on the nucleotide sequences (the calculation of the principal components of the Jaccard similarity matrix) are too computationally intensive to be carried out for all available sequences on GISAID. For this reason, we down-sample each dataset by drawing an unbiased sample of size 10,000 without replacement.

Using the alignment tool MAFFT (Katoh et al, 2002) and the official SARS-CoV-2 reference sequence (available on GISAID under the accession number EPI_ISL_402124), we align all n sequences to the reference genome. We employed MAFFT with the *keeplength* option in order to obtain a well-defined window (of length L=29891 base pairs) for comparison of all sequences. All other parameters of MAFFT were kept at their default values.

### 2.2 Assessing the similarity of nucleotide sequences

We next convert all sequences into a binary Hamming matrix X ∈ B^n x L^ (where B={0,1} is the set of binary numbers) as follows. We compare the reference genome to each aligned nucleotide sequence, and set X_ij_=1 if the sequence with number i differs at position j from the reference sequence. Otherwise, we set X_ij_=0. Here, the number of rows of X is set to the number of nucleotide sequences, and L=29891 is the number of base pairs in the comparison window. The row sums of X correspond to the Hamming distance of each nucleotide sequence to the reference genome. This methodology has already been used in the literature (Hahn et al., 2020a,b,e,f).

We employ the Jaccard similarity measure (Jaccard, 1901; Prokopenko et al, 2016; Schlauch et al, 2017) to assess the similarity of all pairs of sequences. To be precise, each entry (i,j) of the Jaccard matrix 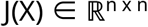 (having n rows and n columns) is a measure of similarity between the binary rows i and j of X. An entry (i,j) of J(X) of zero encodes that the two genomes do not share any deviations from the reference genome, while an entry of one encodes equality of rows I and j of X. We employ the R-package “locStra”, available on CRAN (Hahn et al, 2020c,d), to compute the Jaccard matrix.

For all figures included in this work, we visualize the Jaccard similarity measures by computing its first two principal components. We plot the first principal component against the second principal component, thus effectively interpreting the entries of the first eigenvector as x-coordinates, and the ones of the second eigenvector as y-coordinates. We color each point according to either a time stamp, according to its cluster membership, or according to whether it is an outlier.

### 2.3 Outlier detection

We are interested in detecting sequences falling into neighborhoods or clusters in which they are classified as outliers (subject to a certain criterion). To be precise, we are interested in sequences falling into neighborhoods consisting of sequences having much older (or newer) time stamps.

We aim to utilize an approach which is not dependent on previously identified clusters. One way to achieve this is to define a local environment of radius eps > 0 around each sequence in a principal component plot (each sequence corresponds to a point in the principal component plot), and to consider all other (that is, similar) sequences falling into that local environment. Comparing the time stamp of the sequence under consideration to the distribution of timestamp in the local environment allows one to define an outlier.

We say that a sequence is an outlier in its local environment if its time stamp is more than f > 0 standard deviations from the mean date in the environment.

### 2.4 Calibration

Our clustering approach depends on two tuning parameters, the radius of the local environment eps, and the factor f that specifies how many standard deviations away from the mean date are needed to define a sequence as an outlier. To calibrate both parameters, we look at the number of outliers which are identified in the data as a function of both eps and f. This results in a typical “elbow” plot, though here in two dimensions (see Figure 2). For small values of f, meaning values close to the mean, many outliers are flagged. As f increases, fewer and fewer outliers are identified. The decrease is usually not linear. Instead, the number of outliers usually drops rapidly at a certain cutoff f before leveling off, thus giving the plot its name. The point at which the plot levels off can be used to determine f. We apply the elbow method to both set the parameter f, as well as the parameter eps.

## 3. Results

We first focus on the newest variant, omicron. Figure 1 shows a plot of the first two principal components of the Jaccard matrix as outlined in Section 2.2. As observed previously (Hahn et al., 2022f) the genomes from GISAID exhibit a particular progression pattern, with older submissions (green) clustering in the middle of the plot, while newer samples (red) cluster at the bottom of the plot. The progression of genomes seems to take place from the early point cloud (green, middle), to genomes with intermediate timestamps (top), to new samples (red, bottom). As also observed in the aforementioned publication, genomes of the omicron strain are most similar to genomes in stemming from early on in the pandemic. This is visible from Figure 1 as omicron samples (triangles) fall into a point cloud of early (green) genomes.

**Figure 1.**
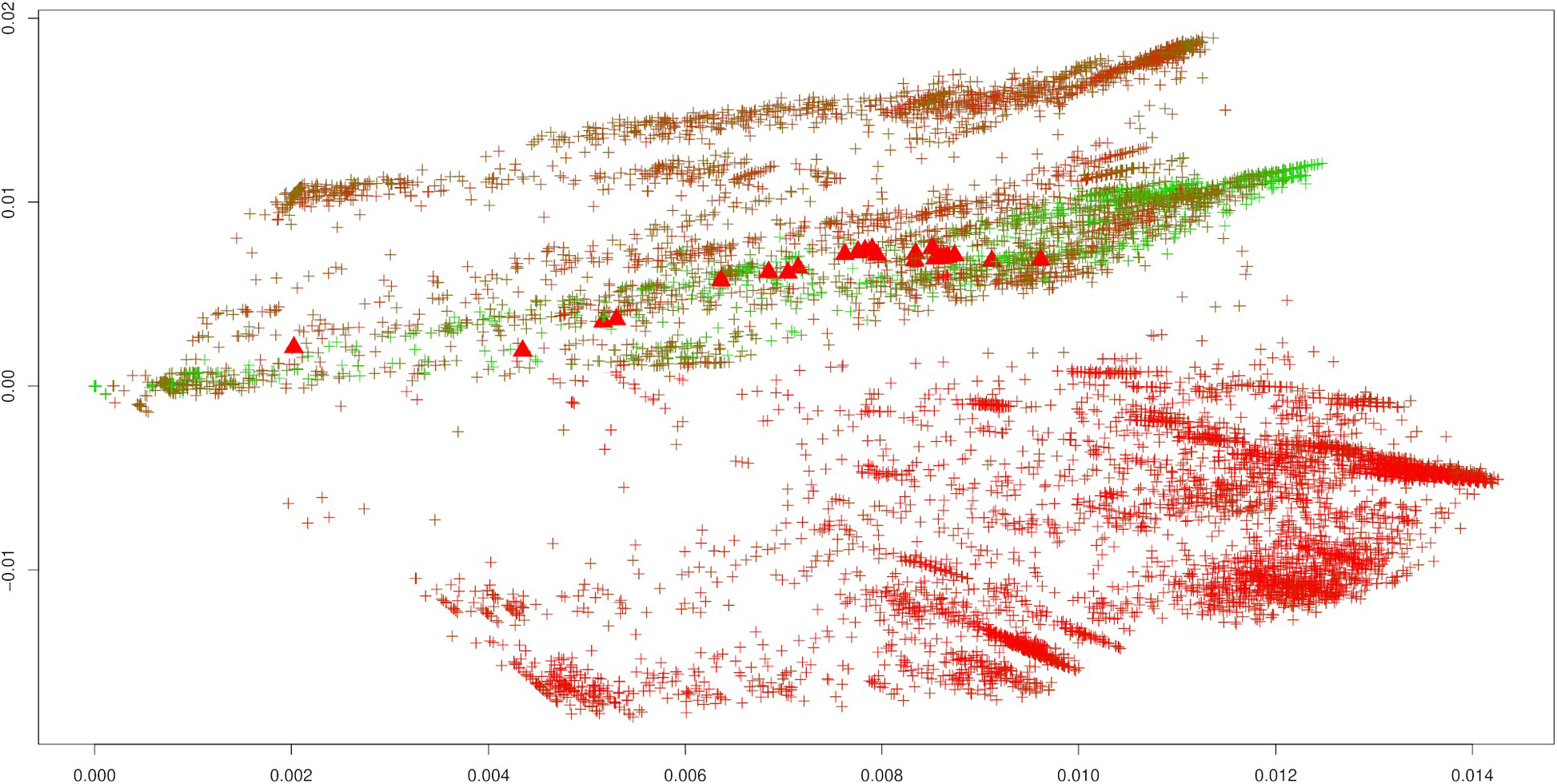
Reference dataset for the omicron variant (see Table 2). First two principal components of the Jaccard matrix, colored by the collection time stamp of each nucleotide sequence. The color scale encodes early (red) to late (green) sequences. Sequences of the omicron variant (see Table 3) are highlighted as triangles.

Interestingly, the observations for Figure 1 are virtually identical with the ones made in Hahn et al. (2022f), even though both experiments are made with independent, and thus entirely different, subsamples without replacement of size 10000 taken from all complete samples available on GISAID.

Before applying the approach of Section 2.3, we calibrate the outlier detection on the omicron data as outlined in Section 2.4. Figure 2 shows the two dimensional elbow plot of the number of flagged outliers as a function of both the radius of the local environment eps and the parameter f. We indeed observe a distinct shape of the decrease in the number of outliers as the parameter f increases, with a sharp decrease at around f=1.2, after which the plot levels off. Interestingly, the algorithm is rather insensitive to the choice of the local environment eps, apart from the case eps=0. We repeated the calibration for the other variants as well. Interestingly, the parameters f=1.2 and eps=1e-1 emerge as consistent choices for all variants. Therefore, we use f=1.2 and eps=1e-1 in the remainder of the section.

**Figure 2.**
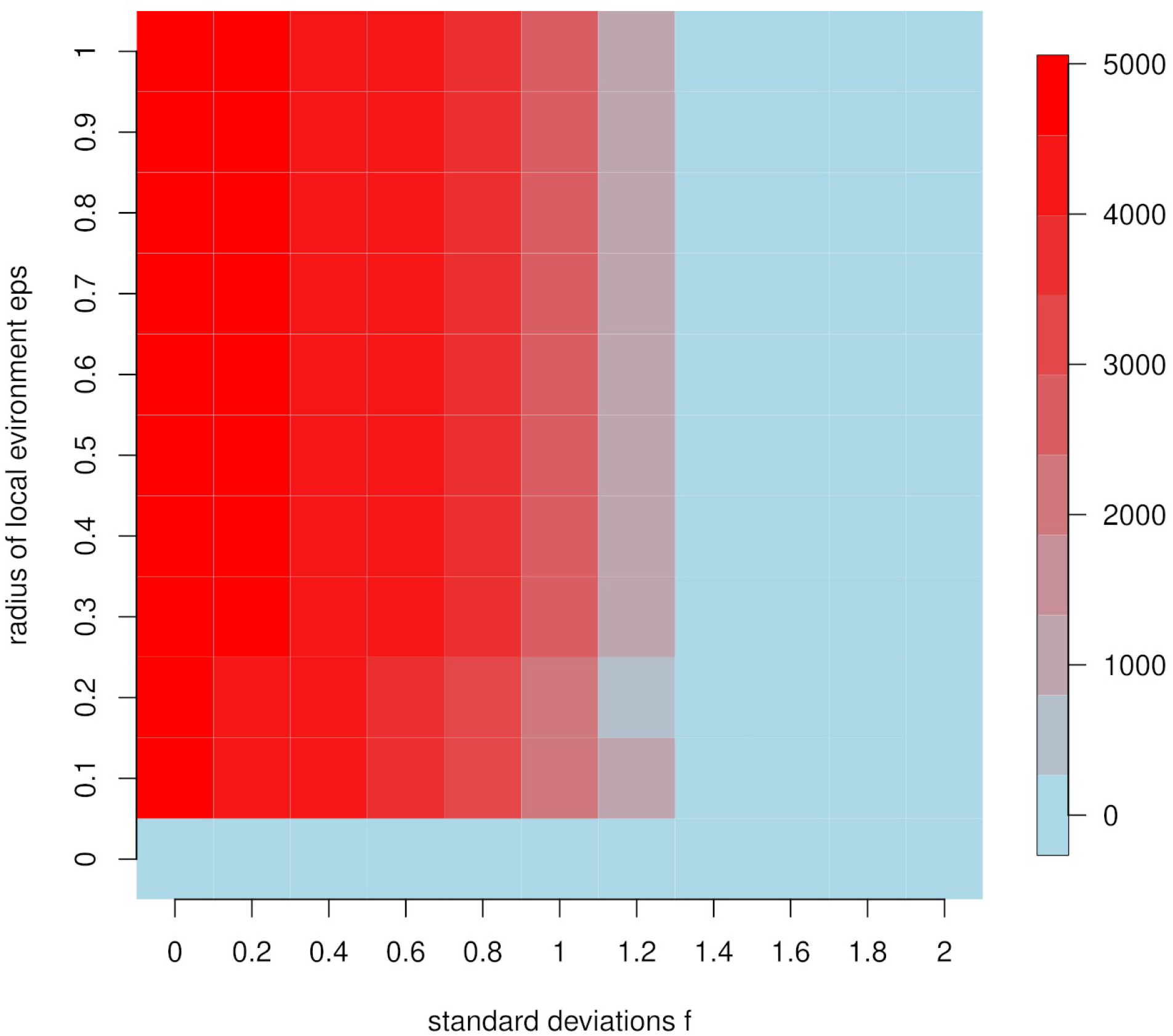
Omicron variant. Heatmap showing the number of outliers (from low, depicted in light blue, to high, depicted in red) as a function of the radius of the local environment eps and the number of standard deviations f.

After calibration, we aim to identify outliers using the local detection approach of Section 2.3. Figure 3 shows the same principal components as Figure 1 for the omicron variant, though this time without any coloring by timestamp. Instead, all points in yellow have the property that they pass the local outlier criterion of Section 2.3, meaning that they are outliers in a local epsilon environment centered around them, subject to the calibration of Section 2.4. As summarized in Table 1, of the 25 omicron genomes included in the dataset, 19 are indeed detected (though with a large false positive rate as there are 276 outliers in Figure 4).

**Figure 3.**
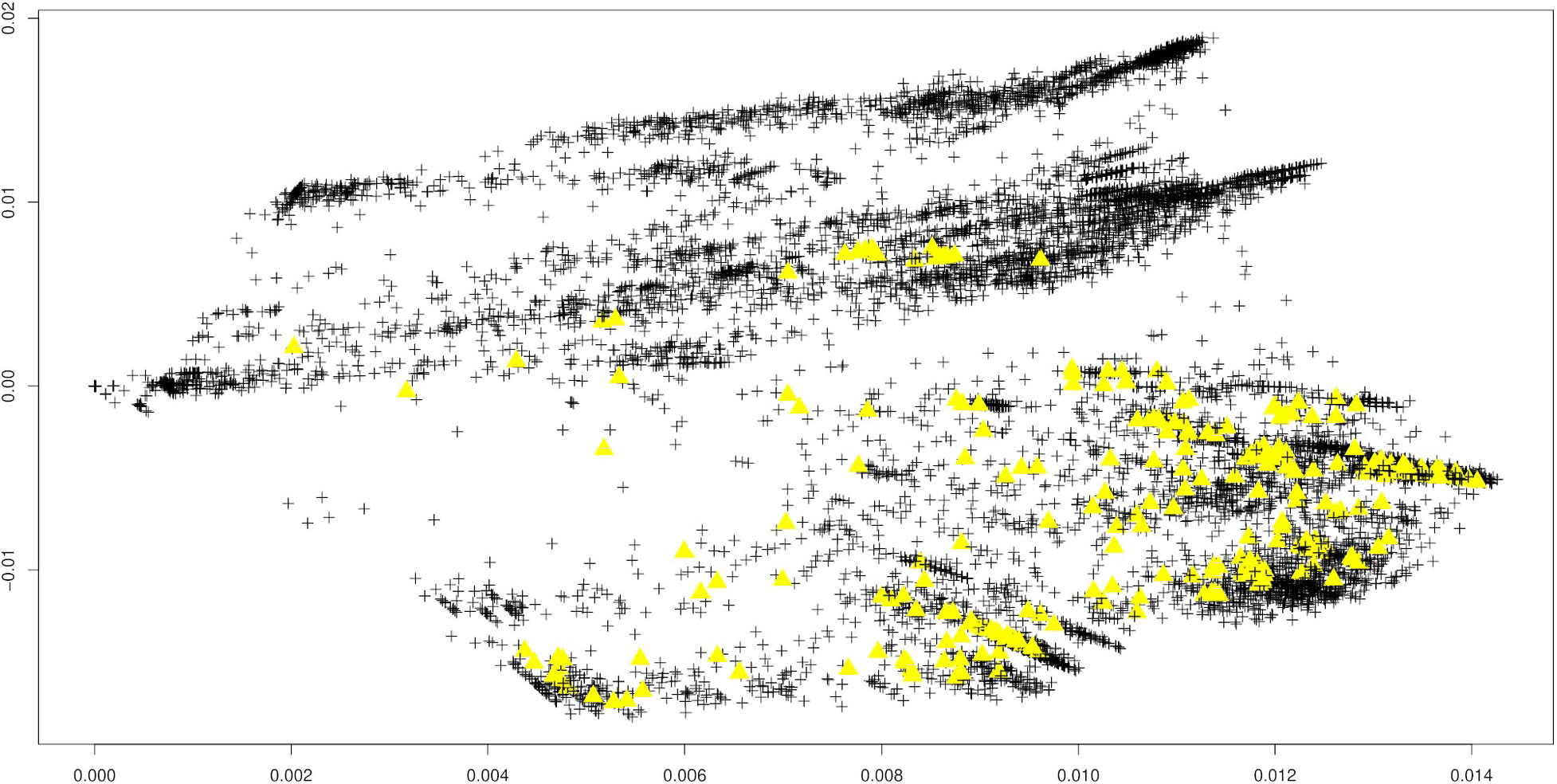
Reference dataset for the omicron variant (see Table 2). First two principal components of the Jaccard matrix with subsequent local outlier detection approach. Parameters eps = 1e-2 (the neighborhood radius) and f=1.5 (the multiplier for the standard deviations). Outliers depicted as yellow triangles.

**Figure 4.**
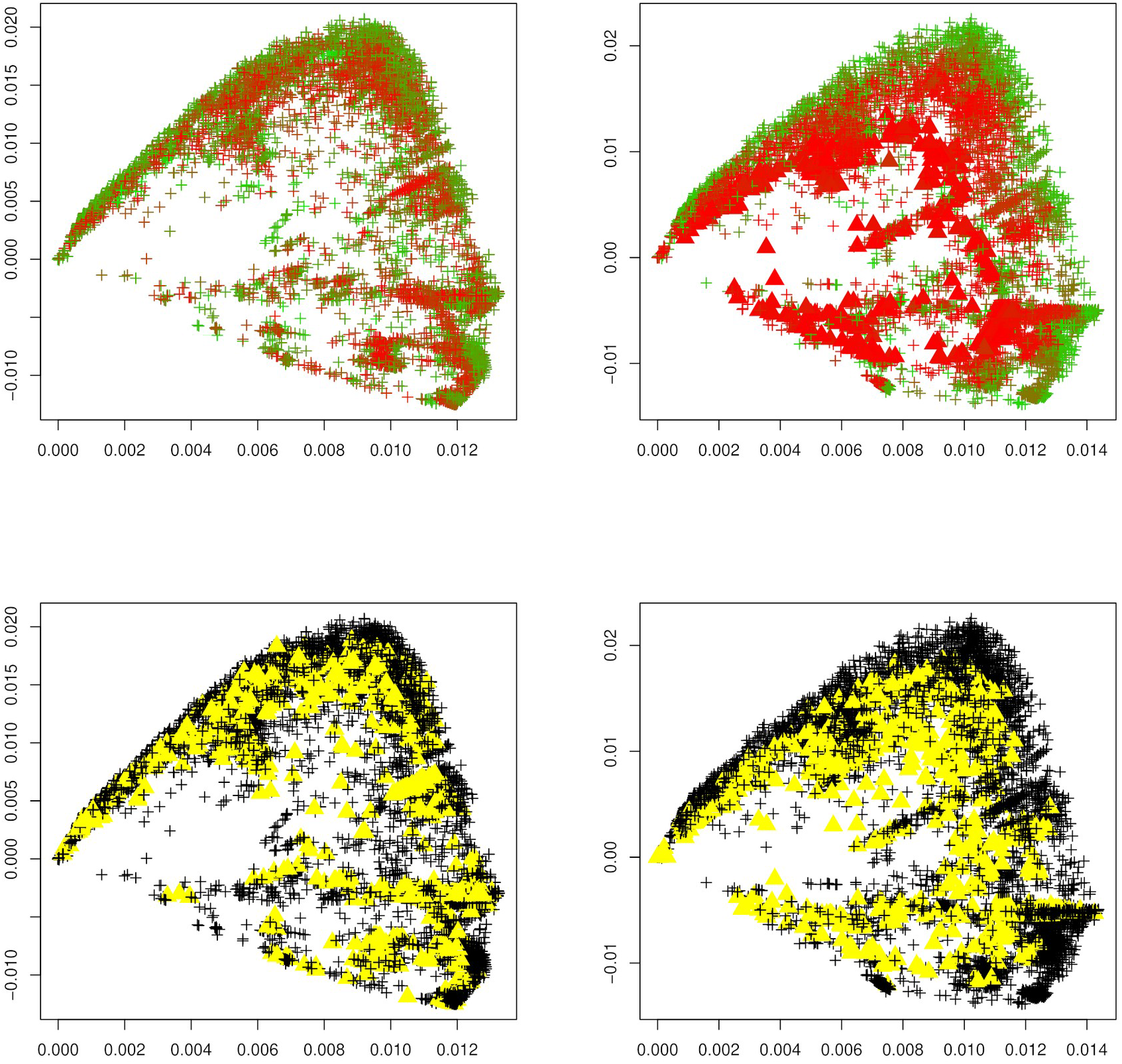
Alpha variant. First two principal components of the Jaccard matrix applied to the reference dataset for the alpha variant (see Table 2) before (top left) and after (top right) the emergence of the alpha variant, where sequences of the alpha variant (see Table 3) are highlighted as triangles. Local outlier detection applied before (bottom left) and after (bottom right) the emergence of the alpha variant, with outliers depicted as yellow triangles.

Interestingly, using the same calibration, many other sequences not belonging to the omicon strain are flagged in Figure 3. These belong to the delta variant of the SARS-CoV-2 virus. In what way these samples differ from the other delta variant samples in Figure 3 remains an important question of future work.

Next, we investigate the question if an increase in the number of outliers can be detected upon the emergence of a new variant. To this end, for each variant under investigation (alpha, beta, delta, gamma, GH, lambda, mu, omicron), we apply the same calibrated outlier detected to first the reference dataset before the emergence of each variant, and after the emergence of each variant. Figures 4-11 show results for all eight variants (alpha, beta, delta, gamma, GH, lambda, mu, omicron). The left column always corresponds to the time period before the emergence of each variant, and the right column corresponds to the time period after the emergence of each variant. The top plots show the first two principal components with highlighted sequences for each variant under consideration, the bottom plots show the local outliers as yellow triangles. We observe that for the beta, delta, GH, and omicron variants the number of detected outliers considerably increases after the emergence of the variant. For the other variants, the change in the number of outliers is less pronounced. For the gamma variant, the number of detected outliers considerably decreases after the emergence of the variant.

**Figure 5.**
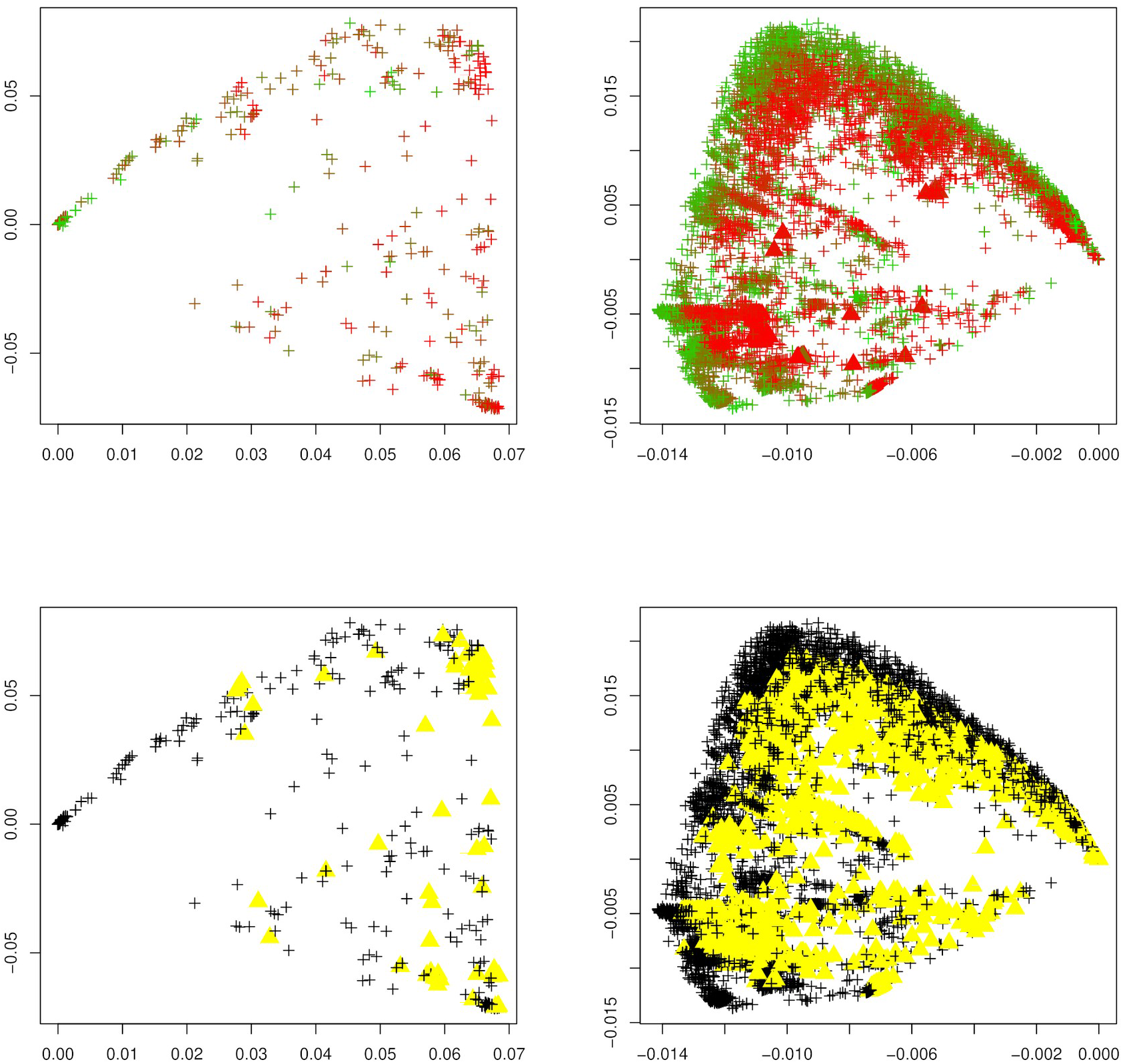
Beta variant. First two principal components of the Jaccard matrix applied to the reference dataset for the beta variant (see Table 2) before (top left) and after (top right) the emergence of the beta variant, where sequences of the beta variant (see Table 3) are highlighted as triangles. Local outlier detection applied before (bottom left) and after (bottom right) the emergence of the beta variant, with outliers depicted as yellow triangles.

**Figure 6.**
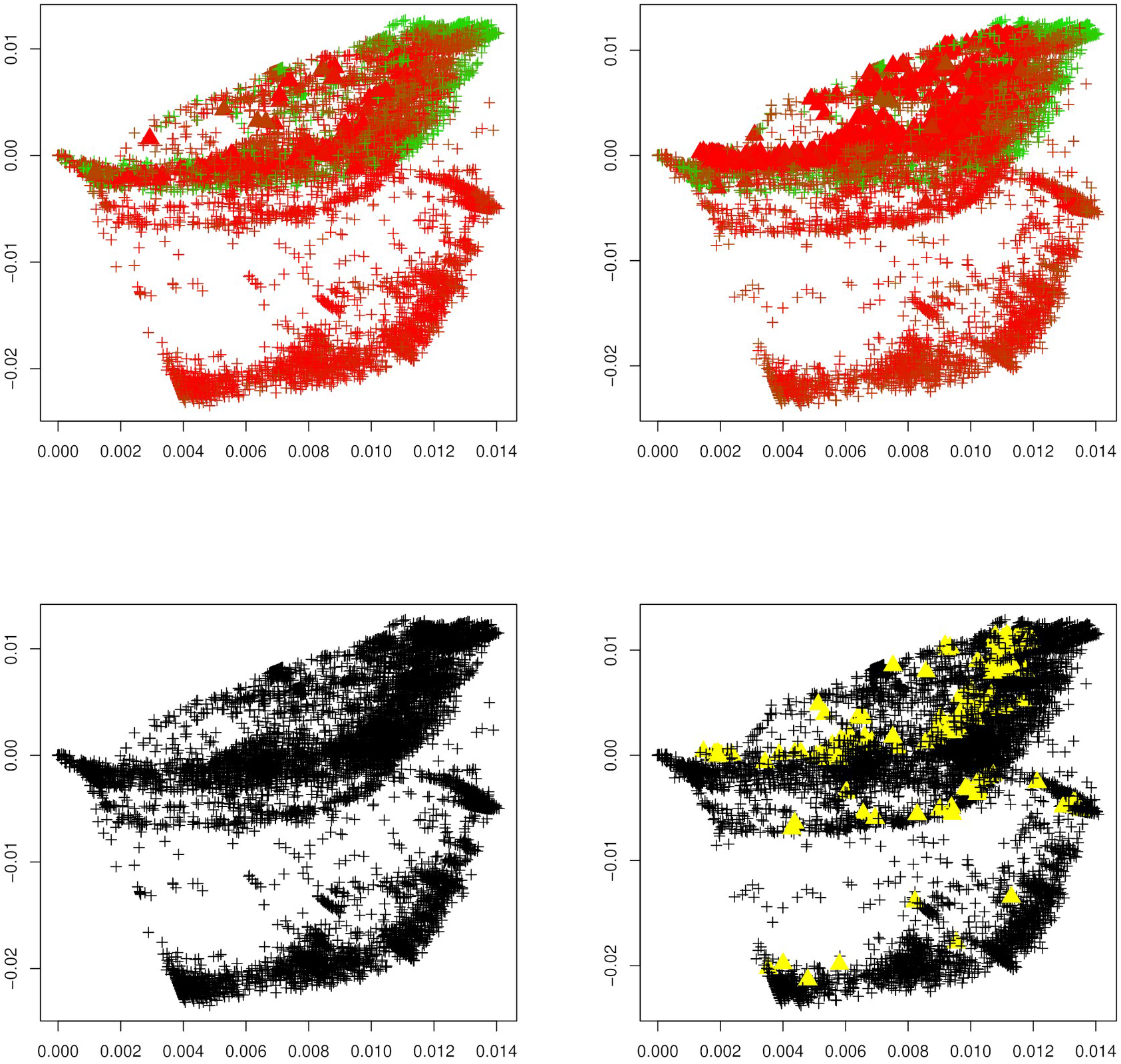
Delta variant. First two principal components of the Jaccard matrix applied to the reference dataset for the delta variant (see Table 2) before (top left) and after (top right) the emergence of the delta variant, where sequences of the delta variant (see Table 3) are highlighted as triangles. Local outlier detection applied before (bottom left) and after (bottom right) the emergence of the delta variant, with outliers depicted as yellow triangles.

**Figure 7.**
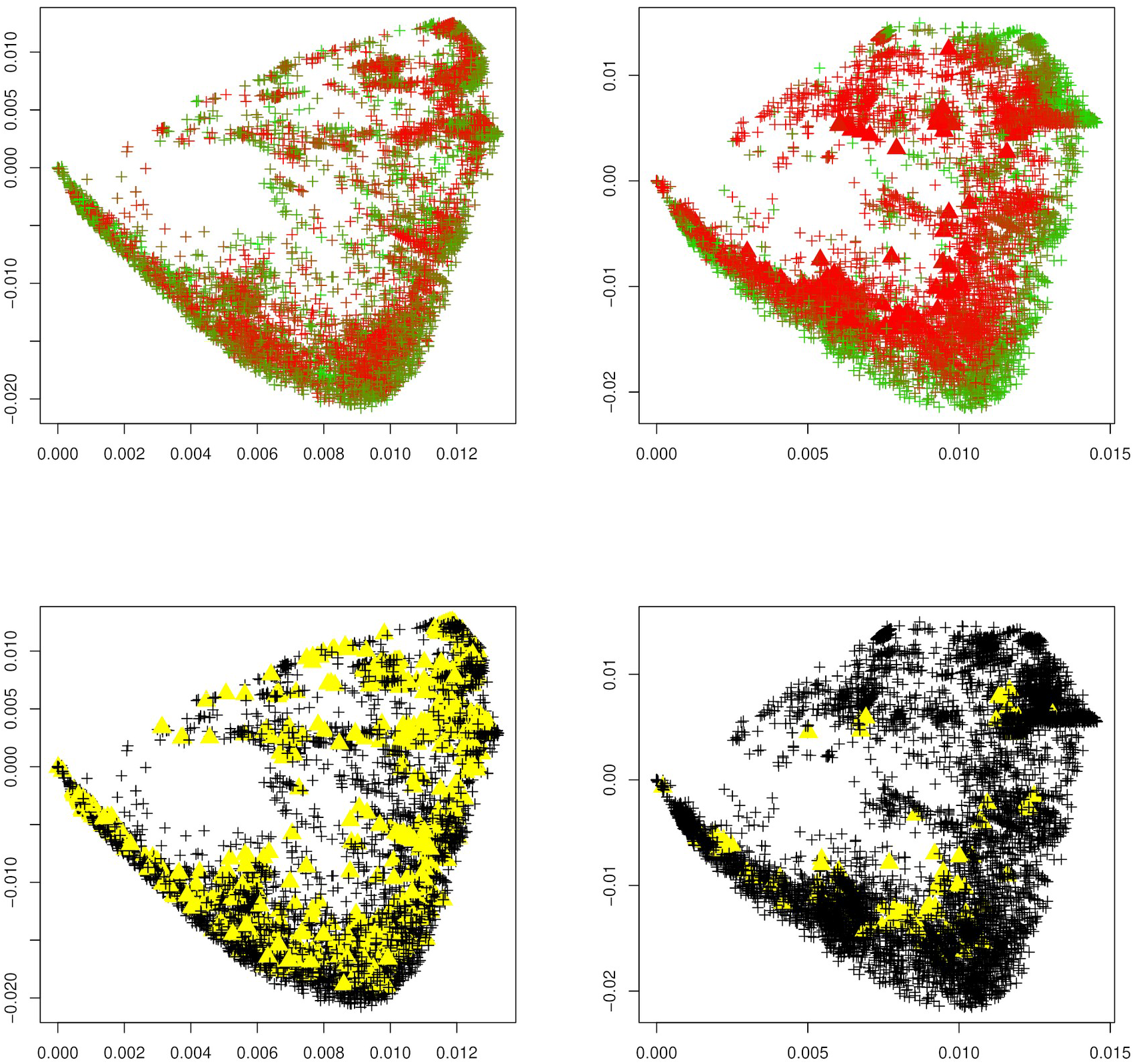
Gamma variant. First two principal components of the Jaccard matrix applied to the reference dataset for the gamma variant (see Table 2) before (top left) and after (top right) the emergence of the gamma variant, where sequences of the gamma variant (see Table 3) are highlighted as triangles (top right). Local outlier detection applied before (bottom left) and after (bottom right) the emergence of the gamma variant, with outliers depicted as yellow triangles.

**Figure 8.**
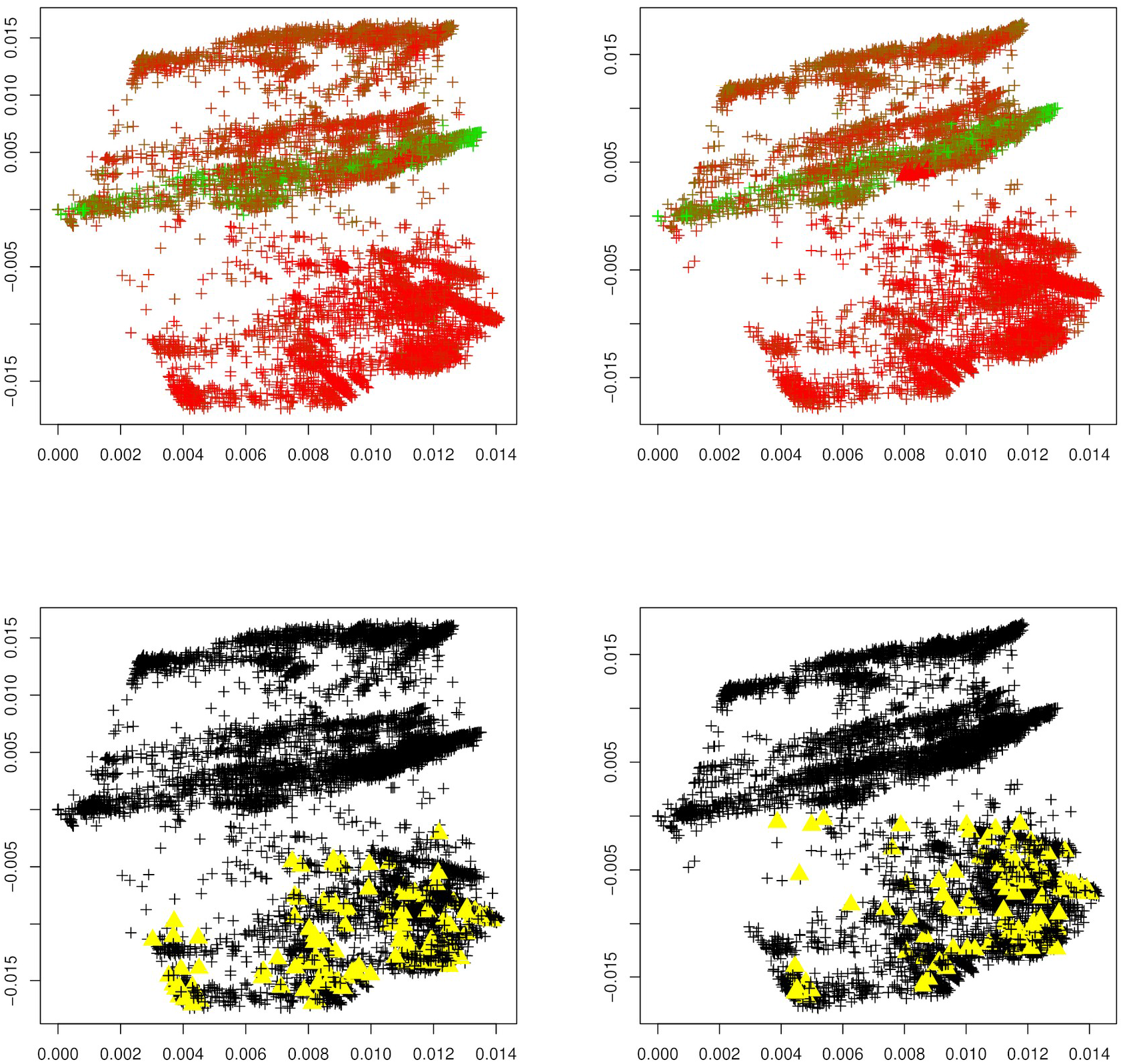
GH variant. First two principal components of the Jaccard matrix applied to the reference dataset for the GH variant (see Table 2) before (top left) and after (top right) the emergence of the GH variant, where sequences of the GH variant (see Table 3) are highlighted as triangles (top right). Local outlier detection applied before (bottom left) and after (bottom right) the emergence of the GH variant, with outliers depicted as yellow triangles.

**Figure 9.**
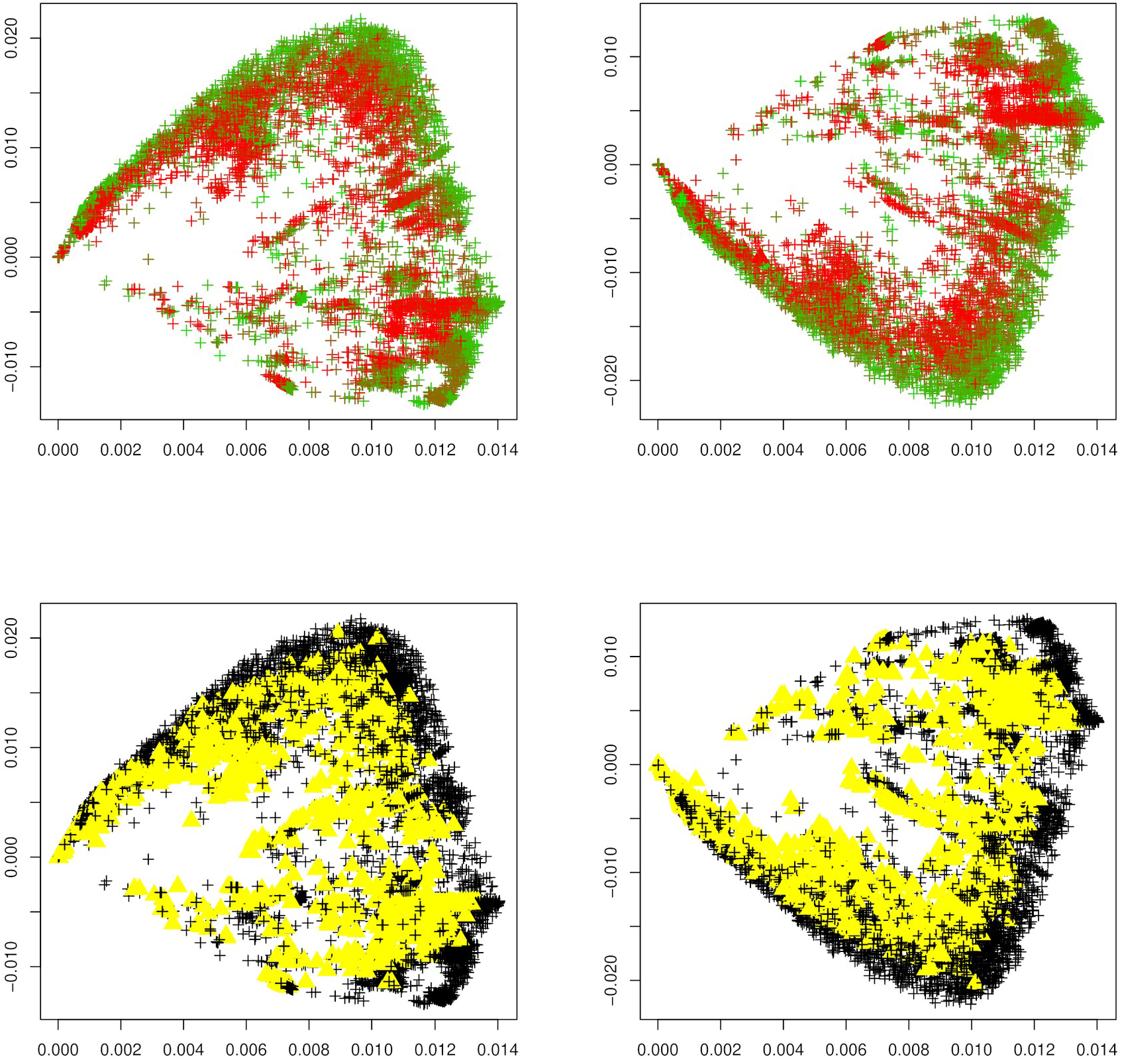
Lambda variant. First two principal components of the Jaccard matrix applied to the reference dataset for the lambda variant (see Table 2) before (top left) and after (top right) the emergence of the lambda variant, where sequences of the lambda variant (see Table 3) are highlighted as triangles (top right). Local outlier detection applied before (bottom left) and after (bottom right) the emergence of the lambda variant, with outliers depicted as yellow triangles.

**Figure 10.**
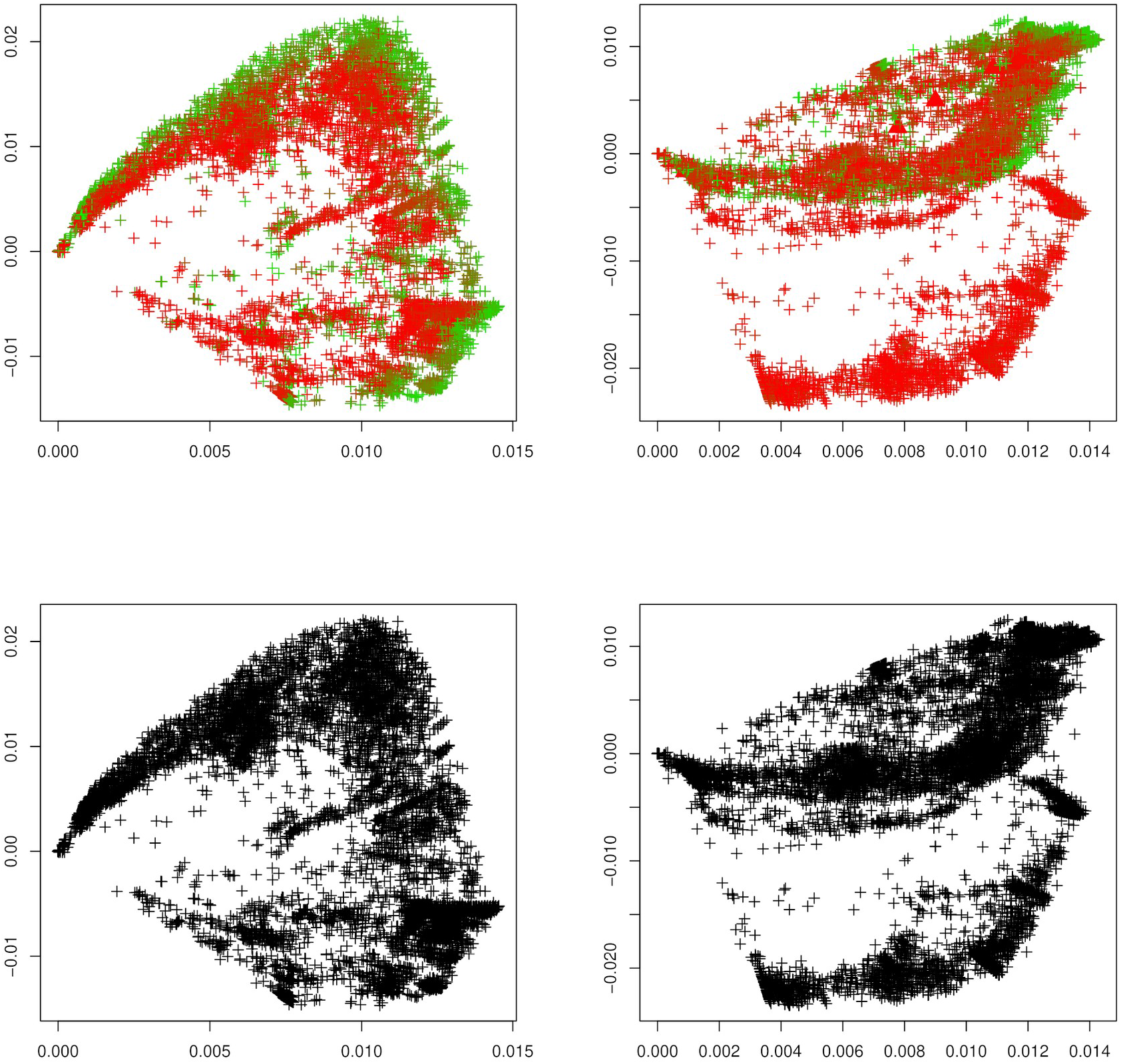
Mu variant. First two principal components of the Jaccard matrix applied to the reference dataset for the mu variant (see Table 2) before (top left) and after (top right) the emergence of the mu variant, where sequences of the mu variant (see Table 3) are highlighted as triangles (top right). Local outlier detection applied before (bottom left) and after (bottom right) the emergence of the mu variant, with outliers depicted as yellow triangles.

**Figure 11.**
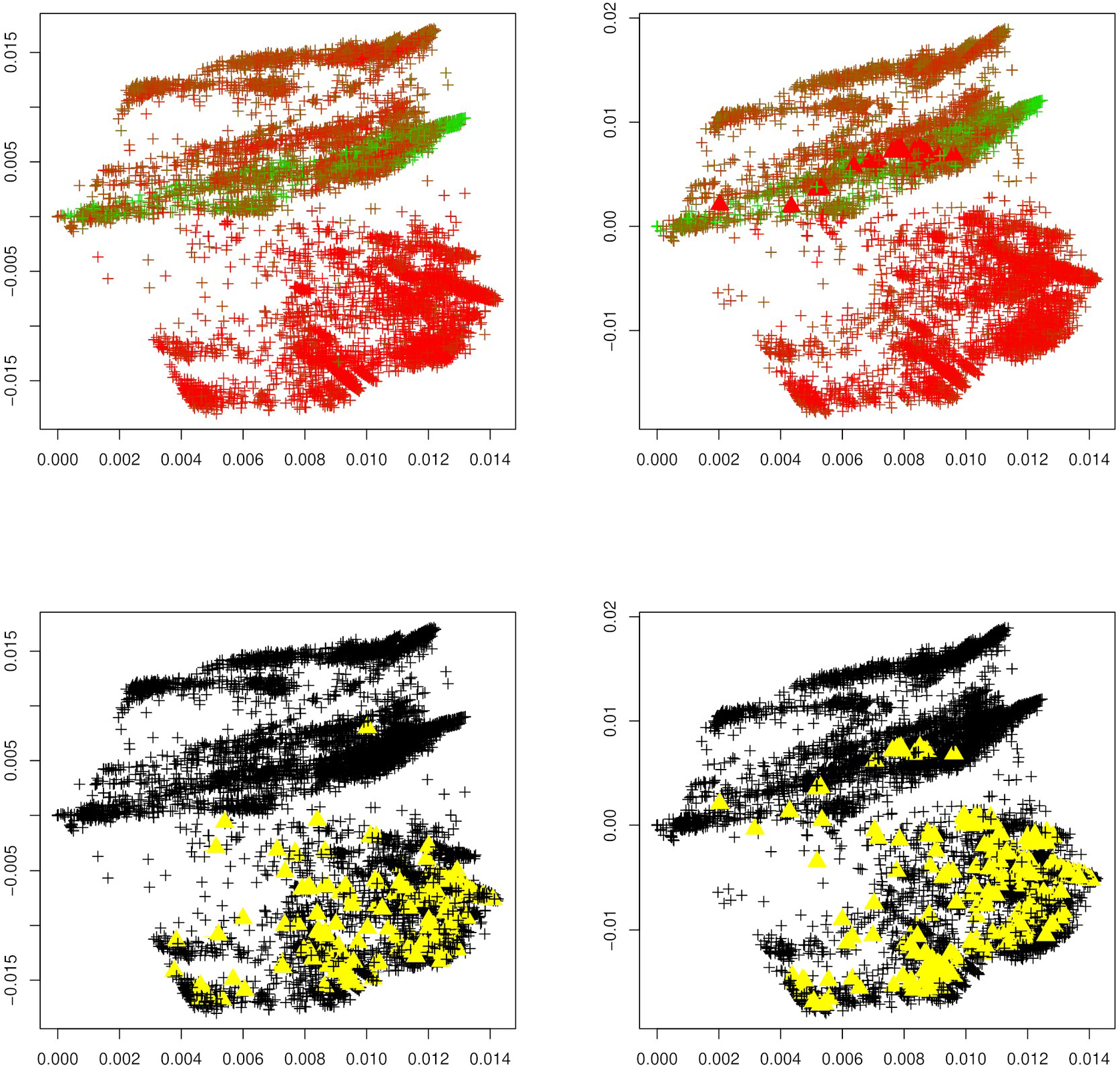
Omicron variant. First two principal components of the Jaccard matrix applied to the reference dataset for the omicron variant (see Table 2) before (top left) and after (top right) the emergence of the omicron variant, where sequences of the omicron variant (see Table 3) are highlighted as triangles (top right). Local outlier detection applied before (bottom left) and after (bottom right) the emergence of the omicron variant, with outliers depicted as yellow triangles.

To concretize results, Table 1 summarizes the total number of detected outliers, the number of detected genomes per variant, and the number of genomes for each variant that is included in the dataset (and that can possibly be detected). We observe that for the common variants beta, delta, GH, and omicron, the detection of the emergence of a new strain is possible. Clearly the biological importance of a new variant cannot be assessed via outlier detection, but the proposed method would have been able to flag these strains as variants of interest.

## 4. Discussion

In this work, we demonstrate that nucleotide sequences of common virus strains/variants can be identified solely based on a statistical outlier criterion in real time. To this end, we prepare two reference datasets, one before and one after the emergence of eight common SARS-CoV-2 variants (alpha, beta, delta, gamma, GH, lambda, mu, omicron) available on the GISAID database, and apply outlier two detection methods to those datasets.

Using the proposed local outlier detection approach, we can identify genomes belonging to the beta, delta, GH, and omicron strain upon emergence of these variants. However, this detection comes at the cost of a larger number of false positives. The nature of those other nucleotide sequences that pass our outlier criteria, and in what way they differ from other sequences of the most common SARS-CoV-2 variants, is an important direction of ongoing research.

Importantly, this research shows that outlier detection might be a useful tool to identify emerging variants in real time as the pandemic progresses, using machine learning techniques and purely statistical methods only.

